# Chemoenzymatic Synthesis Planning Guided by Reaction Type Score

**DOI:** 10.1101/2024.10.25.620365

**Authors:** Hongxiang Li, Xuan Liu, Guangde Jiang, Huimin Zhao

**Affiliations:** NSF Molecular Maker Lab Institute, University of Illinois at Urbana-Champaign, Urbana, IL 61801, USA; Department of Chemical and Biomolecular Engineering, University of Illinois at Urbana-Champaign, Urbana, IL 61801, USA; Carl Woese Institute for Genomic Biology, University of Illinois at Urbana-Champaign, Urbana, IL 61801, USA; Department of Chemistry, University of Illinois at Urbana-Champaign, Urbana, IL 61801, USA; DOE Center for Advanced Bioenergy and Bioproducts Innovation, University of Illinois at Urbana-Champaign, Urbana, IL 61801, USA

**Keywords:** computer-aided synthesis planning, chemoenzymatic synthesis, text-based convolutional neural network, drug synthesis, biocatalysis

## Abstract

Thanks to the growing interests in computer-aided synthesis planning (CASP), a wide variety of retrosynthesis and retrobiosynthesis tools have been developed in the past decades. However, synthesis planning tools for multi-step chemoenzymatic reactions are still rare despite the widespread use of enzymatic reactions in chemical synthesis. Herein we report a reaction type score (RTscore)-guided chemoenzymatic synthesis planning (RTS-CESP) strategy. Briefly, the RTscore is trained using a text-based convolutional neural network (TextCNN) to distinguish synthesis reactions from decomposition reactions and evaluate synthesis efficiency. Once multiple chemical synthesis routes are generated by a retrosynthesis tool for a target molecule, RTscore is used to rank them and find the step(s) that can be replaced by enzymatic reactions to improve synthesis efficiency. As proof of concept, RTS-CESP was applied to 10 molecules with known chemoenzymatic synthesis routes in literature and was able to predict all of them with six being the top-ranked routes. Moreover, RTS-CESP was employed for 1000 molecules in the boutique database and was able to predict the chemoenzymatic synthesis routes for 554 molecules, outperforming ASKCOS, a state-of-the-art chemoenzymatic synthesis planning tool. Finally, RTS-CESP was used to design a new chemoenzymatic synthesis route for the FDA-approved drug Alclofenac, which was shorter than literature-reported route and have been experimentally validated.

## Introduction

Organic synthesis has been renowned for its long history and regarded as the primary choice in synthesizing target molecules for drugs^1,2^, materials^3^ and natural products^4^ for years. However, with the rapid development of biocatalysis and directed evolution in the past decade^5^, various new transformations were imported into the traditional organic synthesis space^6-8^. Owing to its high stereoselectivity and regioselectivity, enzymatic catalysis has been increasingly employed to replace chemical reactions in synthesis routes and selectively produce the desired products^9,10^. Since one enzyme might replace multiple chemical reactions, the chemoenzymatic synthesis route can be shorter and with higher yields. For instance, in the synthesis of sitagliptin^11^, the overall hybrid synthesis route was shortened by three steps compared to the original chemical synthesis route and the enantiomeric excess was also increased with enzyme catalysis. Moreover, biocatalysis uses mild conditions and avoids toxic reagents or high pressure, facilitating green chemistry^12,13^, while the one-pot enzymatic cascades help reduce the purification between reactions^14-17^. Not surprisingly, chemoenzymatic synthesis has been applied to multiple practically important small molecules^18-21^.

Other than human efforts and expertise in designing synthesis routes, the computer-aided synthesis planning (CASP) tools have been explored for both organic and enzymatic synthesis^22-25^. CASP is a process of breaking down target molecules step by step, using either machine learning (ML) or rule-based methods, until reaching commercially available molecules. The rule-based CASP tools apply well-defined reaction rules (also called templates) to each target molecule and generate corresponding precursors (ML algorithms can be used for selecting templates) while the ML-based CASP tools predict the precursors using algorithms trained on reaction databases. Synthia^25^ is a commercial retrosynthesis tool using chemist curated rules, while Aizynthfinder^26^ uses abundant extracted rules from the Reaxys database^27^ and employs Monte Carlo tree search (MCTS) to save calculation time. For retrobiosynthesis, novoStoic^28^ focuses on enzymes in metabolic engineering and aims to synthesize target molecules from metabolites, while RetroBioCat^29^ specializes in enzymatic cascades for *in vitro* synthesis and uses curated reaction rules for prediction. As the first ML-based retrobiosynthesis tool, the IBM’s RXN4Chemistry^30^ adapts transformer with ECREACT database^30^ and can potentially predict novel reactions.

However, CASP tools for chemoenzymatic synthesis planning remain rare despite that many chemoenzymatic synthesis routes have been reported. To the best of our knowledge, there are only three chemoenzymatic synthesis planning tools reported so far. In 2022, Coley and co-workers designed ASKCOS (hybrid version) ^31^, which extracts templates from the Reaxys (chemical) and BKMS (biological) databases^32^, and then uses two prioritizers to rank those templates in each step and expand the search tree. Since the prioritizers are based on reaction templates, this tool is limited to only rule-based methods. Moreover, because of their step by step searching algorithm, there could be multiple transitions between chemical and biological reactions in their predicted routes, increasing difficulties in experiments. Later, Jensen and co-workers^33^ developed an alternative tool in which they firstly used ASKCOS to generate chemical synthesis routes and then identified enzymes to carry out the same transformation for every step using templates in RetroBioCat. This tool did not aim to predict new enzymatic transformations, and it is time-consuming to exhaustively search for alternative enzymatic reactions to replace every chemical step in the original routes. Recently, Wu and co-workers developed BioNavi^34^ which only used ML-based methods, and their biological reaction predictor was specifically trained for natural product synthesis. Notably, in the first two studies, no experimental validation was conducted for the newly proposed reactions, and only partial synthesis routes were validated in BioNavi.

In this work, we have developed a reaction type score (RTscore)-guided chemoenzymatic synthesis planning (RTS-CESP) strategy (Figure 1). The existing retrosynthesis and retrobiosynthesis tools (e.g., Aizynthfinder^2629^, RXN4Chemistry^30,35^) were employed to identify chemical and biological reactions, which were then integrated by a custom-designed automatic searching algorithm to generate chemoenzymatic synthesis routes for the target molecules. To start with, an RTscore is designed by training a text-based convolutional neural network (TextCNN) to distinguish synthesis reactions from decomposition reactions, which could achieve an F1 score of 0.971 using the USPTO dataset^36^. For each target molecule, the chemical synthesis route is generated first, and then RTscore is applied to every step to predict reaction type and determine which steps should be replaced by enzymatic reactions. As proof of concept, we used this tool to predict the synthesis routes for 10 molecules with known chemoenzymatic synthesis routes in literature and found all these reported chemoenzymatic synthesis routes were successfully predicted and six of them were ranked as the most preferred routes. Furthermore, a validation set with 1000 molecules was adopted to evaluate the performance of this tool on a large scale. Among the 1000 molecules from the boutique database^37^, our tool could predict chemoenzymatic synthesis routes for 554 molecules, versus 493 molecules by the state-of-art tool named ASKCOS, and we proposed shorter pathways for 30.2% molecules that were found by both tools. In addition to reproducing literature results, we predicted a novel shorter pathway for an FDA-approved drug Alclofenac^38^ and experimentally validated the synthesis route with 69% yield.

**Figure 1.**
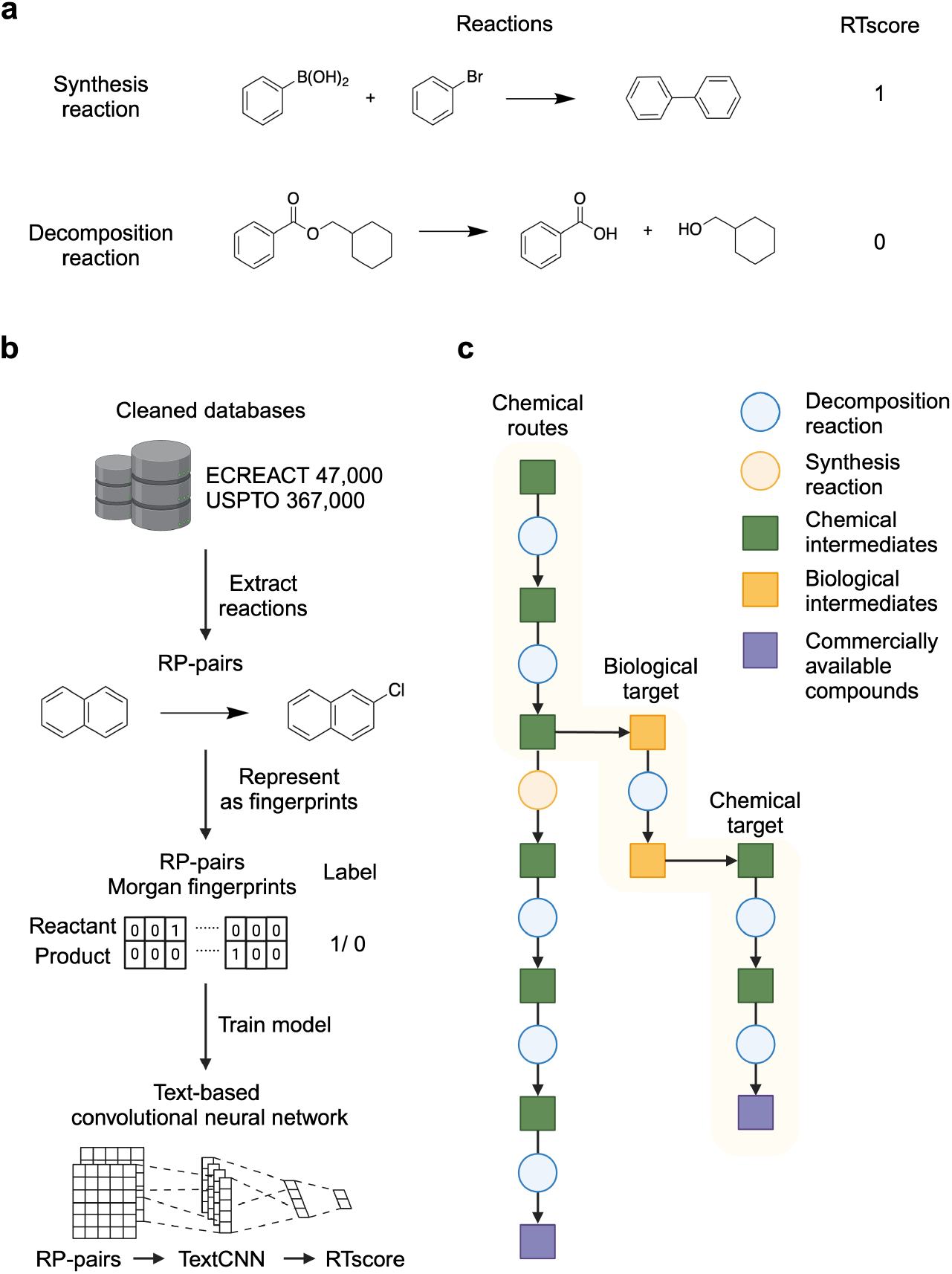
RTscore training process and searching algorithm. **a**, Examples of synthesis reactions and decomposition reactions. In the process of breaking down the target molecules in a retrosynthesis manner, the decomposition reactions are preferred. **b**, Procedure for training the RTscore model. The USPTO and ECREACT databases were first cleaned to remove cofactors in reactions, then the major reactant-product pairs (RP-pairs) were extracted and represented as Morgan fingerprints with labels 1 for a synthesis reaction and 0 for a decomposition reaction separately. With a text-based convolutional neural network, the model predicts RTscore with the input of RP-pairs. **c**, Hybrid synthesis search algorithm. The chemical synthesis route is generated for the target molecule at first and RTscore_(USPTO)_ is employed to find the reaction that does not break down the target molecule effectively and search the product molecule in that reaction with retrobiosynthesis tools, then the biological precursor is searched again with the chemical method to form the chemoenzymatic synthesis routes.

## Results

### RTscore for ranking synthesis routes and guiding hybrid synthesis

Retrosynthesis tools are used to break down target molecules into commercially available compounds through multiple steps. To improve synthesis efficiency, each step should ideally decompose the target molecules into simpler molecules rather than synthesizing more complex products (Figure 1a). The synthesis efficiency of each reaction can be evaluated based on several factors, including changes in molecular weight and atom numbers of the reactants and products, the introduction of chirality, and the number of references supporting the reaction. We processed the database to retain reactions where the product has a larger molecular weight or atom number than the reactant. Then we defined reactions as recorded in the database (the reaction from reactant to product) as synthesis reactions, and the reversed reaction (from product to reactant) as decomposition reactions. We designed an RTscore using a text-based convolutional neural network (TextCNN) to distinguish these two reaction types, while also learning chirality changes and synthesis preferences from the database (Supplementary Figure 1). RTscore was applied to each step of the synthesis routes. After summing the scores of all reactions, the entire route can be scored and ranked. Additionally, in each route, the reaction with the lowest score is replaced by an alternative biological step to generate hybrid synthesis routes. Finally, hybrid routes with fewer steps and faster access to commercially available materials are selected as the top options.

Since chemical and biological reactions occupy different reaction spaces, we trained two separate scores using the USPTO (chemical) and ECREACT (biological) databases, forming RTscore_(USPTO)_ and RTscore_(ECREACT),_ respectively (Figure 1b). Our training input comprised the extracted reactant-product pairs (RP-pairs) from both databases, processed as follows. First, common co-factors were removed to avoid interference (Supplementary Figure 2), as our goal was to rank the routes based on the transformation between the major reactant and product rather than predicting a complete reaction. Next, since all the reactions in both databases contain only a single product, it was regarded as the major product. For reactions with multiple reactants, we calculated the fingerprint similarity of all reactants to the major product and the reactant with the highest similarity score to the major product was selected as the major reactant. In addition, all SMILES strings in the dataset were canonicalized to ensure a consistent representation of molecular structures, and we deduplicated the dataset to remove any redundant entries. Finally, we retained the reactions where the major product had a larger molecular weight or more atoms than the major reactant and discarded the others. The retained reactions were labeled as synthesis reactions, while the reversed reactions labeled as decomposition reactions. This systematic approach allowed us to construct a comprehensive dataset containing 367,000 chemical and 47,000 enzymatic RP-pairs. The differentiation of reaction type was defined as a classification problem and the dataset was split into training, validation, and test sets (8:1:1). A performance comparison of four models— Random Forest (RF)^39^, Support Vector Machine (SVM)^40^, Feedforward Neural Network (FNN)^41^, and Text-based Convolutional Neural Network (TextCNN)^42,43^—was conducted using the ECREACT database. The models were evaluated using both the F1 score and the Matthews Correlation Coefficient (MCC), with the TextCNN model achieving the highest values for both metrics (Supplementary Figure 3). Therefore, the TextCNN model was selected for further study. A two-layer TextCNN was employed, utilizing ReLU (Rectified Linear Unit) activation functions after each convolutional and fully connected layer, except for the final layer which uses a sigmoid activation function. In addition, we evaluated the model’s performance using both binary and count-based fingerprints, with the count-based fingerprint achieving higher F1 and MCC scores (Supplementary Figure 4). To further improve the model, we performed hyperparameter tuning using a grid search approach with 5-fold cross-validation. The hyperparameter grid included output channels of 2, 3, and 4 for the first and second convolutional layers and hidden sizes of 256, 512, and 1024 for the fully connected layers that follow the convolutional layers. The heatmaps for F1 scores on the biological and chemical datasets illustrate the performance of different hyperparameter combinations. For the biological dataset ECREACT, the best F1 score of 0.921 was achieved with a hidden size of 512 and output channels of 4 (Supplementary Figure 5). For the chemical dataset USPTO, the highest F1 score of 0.971 was obtained with a hidden size of 256 and output channels of 3 and 4 for the first and second convolutional layer separately (Supplementary Figure 6). To further assess the model’s robustness, we evaluated its performance using 10 different random seeds, which demonstrated stability across runs with a standard deviation of 0.0034 for the ECREACT database and 0.00089 for the USPTO database, confirming the model’s reliability under different initializations (Supplementary Figure 7). After that, we evaluated RTscore_(ECREACT)_ using the BKMS database^32^, an external enzymatic database. We first deduplicated the reactions with the ECREACT database and selected the major RP-pairs, then RTscore_(ECREACT)_ was used to predict reaction type and reached 75% accuracy, while the SCScore^44^ reached only 64% accuracy (Supplementary Figure 8).

### Multi-step searching algorithm

To achieve effective chemoenzymatic synthesis by integrating retrosynthesis and retrobiosynthesis tools, we designed an automatic hybrid synthesis search algorithm (Figure 1c). In this algorithm, a target molecule is first input into a retrosynthesis tool, and RTscore_(USPTO)_ is used to rank the predicted routes. Since RTscore_(USPTO)_ can be calculated for each reaction, the reaction with the worst score in each route is selected for improvement. The product molecule in that reaction is then input into retrobiosynthesis tools to find alternative enzymatic reactions. All available enzymatic templates are explored due to their limited number. RTscore_(ECREACT)_ is then applied to rank the enzymatic reactions and identify intermediates suitable for chemical synthesis. Finally, the hybrid synthesis routes are collected, and the RTscore for each reaction in the entire route is added up to rank all the predicted routes. The interface between the different tools is fully programmatic. Once a target molecule is input, the algorithm automatically generates a list of chemoenzymatic synthesis routes with ranking.

### Validation of RTS-CESP using a dataset containing known hybrid synthesis routes

To determine the prediction accuracy of RTS-CESP, we sought to use the hybrid synthesis routes reported in literature as a test case. We chose 10 target molecules and discovered the corresponding synthesis reactions in literature from a chemoenzymatic synthesis routes dataset^45-62^. The retrosynthesis tools used in this task were RXN4Chemistry (chemical) and RXN4Chemistry (enzymatic), while RTscore was used to guide the search and rank all the predicted chemoenzymatic synthesis routes. As shown in Figure 2, all the reported chemoenzymatic synthesis routes for these 10 molecules were identified and six of them were ranked as the top by RTscore, indicating that RTscore can be used to prioritize the chemoenzymatic synthesis routes and help researchers choose the most efficient routes.

**Figure 2.**
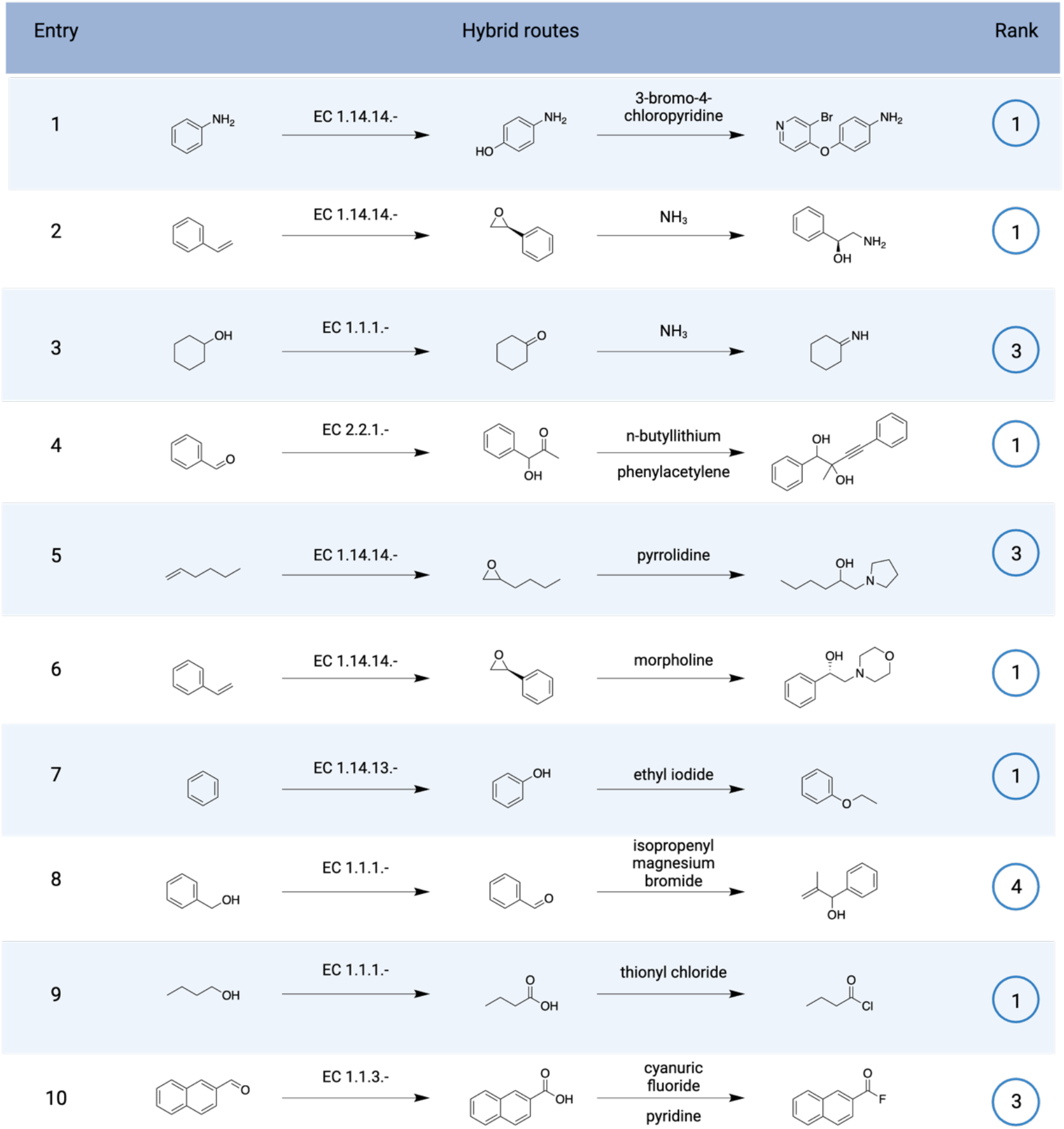
Validation with literature-reported hybrid synthesis routes. For 10 selected target molecules, we searched for literature reactions to build a hybrid synthesis route dataset in which each route contains both chemical and enzymatic reactions. All the hybrid synthesis routes were successfully reproduced by our tool and when ranked with RTscore, 6 out of 10 routes were ranked as the highest among all the predicted routes for a target molecule.

### Large-scale validation and benchmark of RTS-CESP with ASKCOS

To evaluate the performance of RTS-CESP on a large scale, we selected 1000 molecules from the boutique database as target molecules and used Aizynthfinder and RetroBioCat for synthesis planning. Aizynthfinder utilized rules extracted from the Reaxys database and performed well in breaking down complex molecules, so it was used to generate the chemical synthesis routes for a target molecule at first. We then applied the biological reaction templates from the RetroBioCat database to intermediates in these chemical synthesis routes to predict enzymatic reactions. For the target molecules that Aizynthfinder failed to generate a complete chemical synthesis route (a route from target to stock molecules), we selected the intermediates in the generated partial route as targets for biological reactions. Finally, the biological precursors were searched with Aizynthfinder again to finish the whole routes and we ranked all the synthesis routes using RTscore.

Using the same calculation time (three minutes) and stock molecules^31^ (i.e. molecules less than $100/g from eMolecules and Sigma-Aldrich) (Supplementary Table 1), RTS-CESP predicted the chemoenzymatic synthesis routes for 554 molecules, while the state-of-the-art tool ASKCOS hybrid predicted the chemoenzymatic synthesis routes for 493 molecules (Figure 3a) (See Supplementary Figure 9 for examples of chemoenzymatic synthesis routes to target molecules identified by our tool but not by ASKCOS). There were 371 molecules that both tools have identified synthesis routes for, among which RTS-CESP predicted shorter synthesis routes for 112 of them (30.2%) (Figure 3b, c). Besides, as a self-benchmark, employing the chemoenzymatic search algorithm could predict synthesis routes for more molecules than using the chemical synthesis algorithm only (Figure 3d). The calculation time and route length varied for different target molecules (Figure 3e, f). Most target molecules were solved within three minutes, and more than half of the routes consisted of one or two steps, showing RTS-CESP’s effectiveness in planning synthesis routes. Moreover, RTS-CESP could find routes with more than eight steps, indicating its competence in breaking down complex molecules.

**Figure 3.**
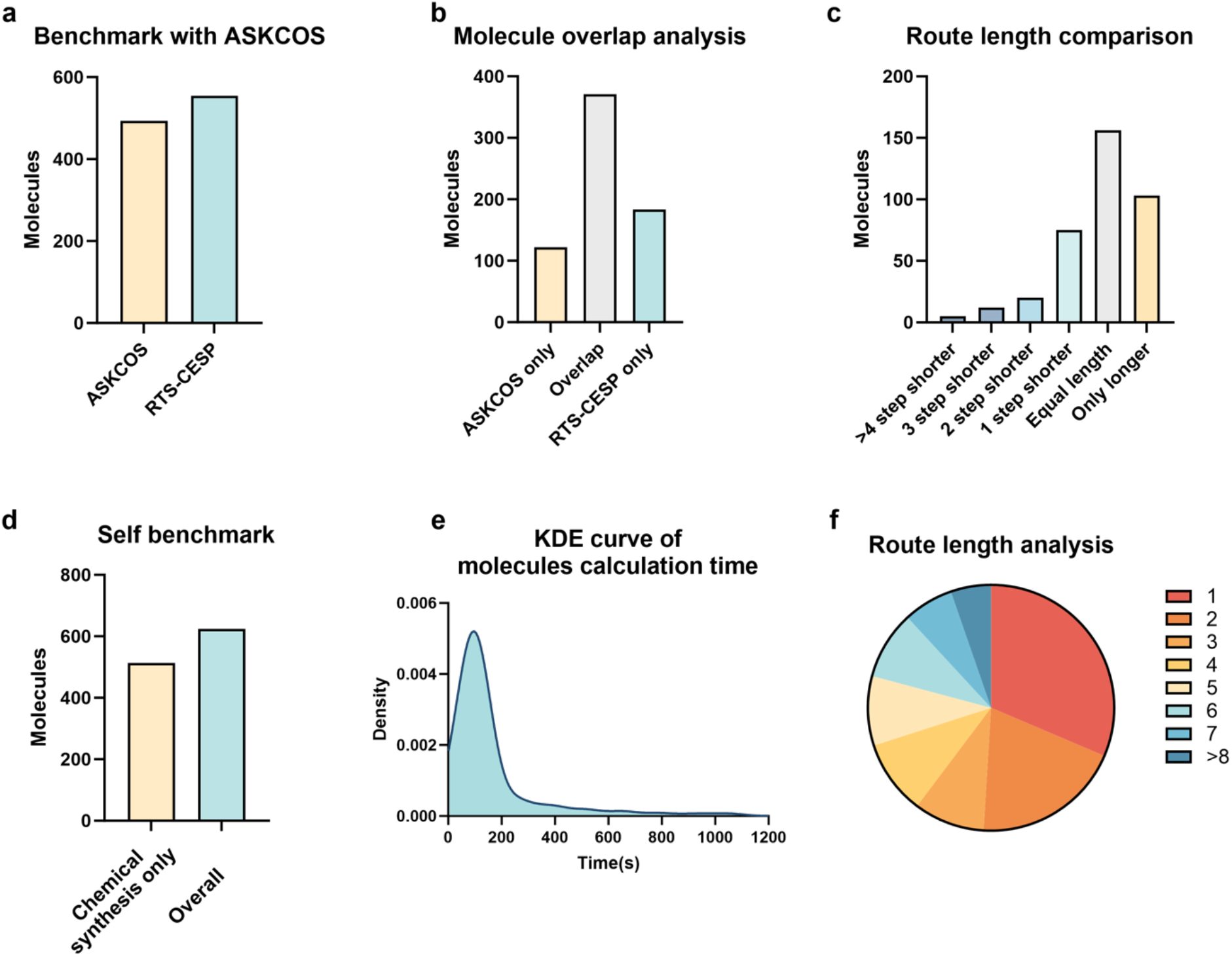
Validation and benchmark study on a large-scale dataset. **a**, Benchmark with ASKCOS on the 1000 target molecules from the boutique database using same calculation time (three minutes) and stock molecules. **b**, Analysis of the molecules whose chemoenzymatic synthesis routes were predicted by RTS-CESP and ASKCOS. **c**, Comparison of the lengths of synthesis routes predicted by RTS-CESP and ASKCOS. RTS-CESP identified more shorter synthesis routes than ASKCOS. **d**, Using the hybrid synthesis search algorithm of RTS-CESP identified synthesis routes for more molecules than using the chemical synthesis search algorithm only. **e**, The kernel density estimation (KDE) curve for molecule calculation time. **f**, Distribution of the lengths of predicted synthesis routes.

**Figure 4.**
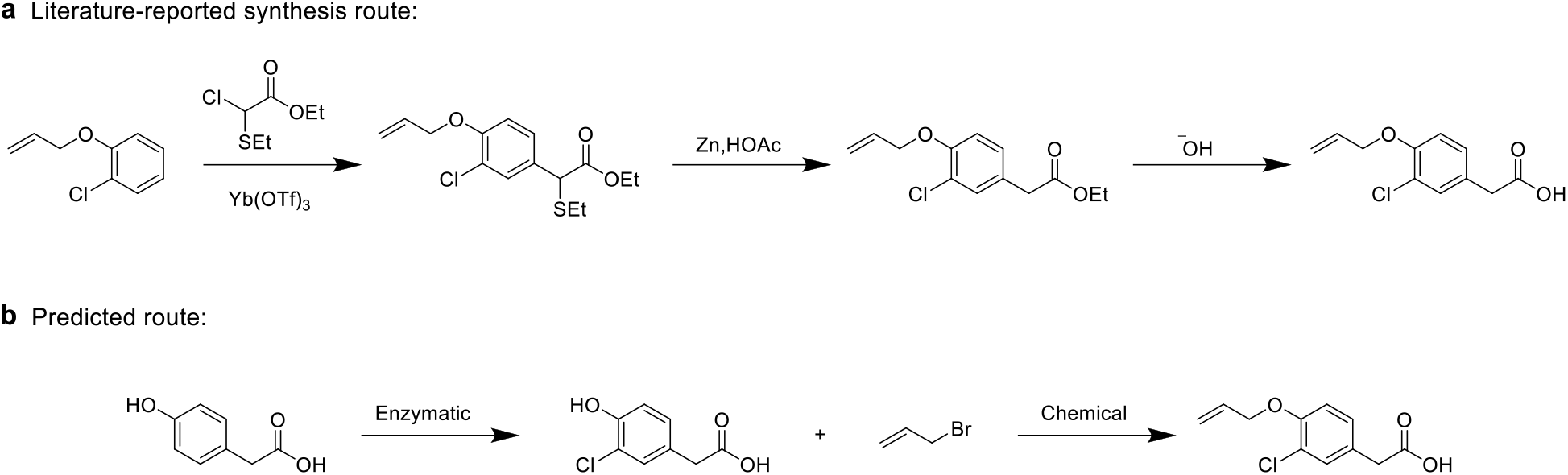
Experimental validation of the predicted synthesis route for an FDA-approved drug. **a**, Literature-reported synthesis routes for Alclofenac. **b**, Our predicted route. Our predicted route is one step shorter than the literature-reported route and starts with a cheaper precursor.

### Experimental validation of the predicted synthesis route for an FDA-approved drug

Alclofenac is an FDA (Food and Drug Administration) approved anti-inflammatory drug. To the best of our knowledge, only chemical methods were used to synthesize this compound and the shortest synthesis route in the literature consisted of three steps^37^. Here, we used RTS-CESP to predict synthesis routes for the target. Aizynthfinder was used to generate the chemical synthesis route first, and templates from RetroBioCat and BKMS databases were used to generate the enzymatic step. A synthesis route with only two steps was found, including one chemical and one enzymatic step and both have not been reported in the literature. Therefore, we sought to experimentally validate this predicted synthesis route.

For the first step (enzymatic halogenation), we tested a previously reported chloroperoxidase^63^ and were able to isolate the target product with 76% yield. For the second step, we performed the reaction in ethanol with base added, generating the final product with 91% yield. Besides that, when we compared the prices for the starting material in our route and the literature-reported route from the same supplier, the compound in literature-reported route is 10 times more expensive than ours.

## Discussion

In this work, we have developed RTS-CESP, a versatile, robust, and reliable chemoenzymatic synthesis planning tool. It starts with the predicted chemical synthesis routes for a target molecule and identifies the steps that do not break down the target molecule efficiently and replaces them with enzymatic reactions, which saves searching time and minimizes transitions between chemical reactions and enzymatic reactions. RTS-CESP has been validated using a small-scale database with 10 known chemoenzymatic synthesis routes, a large-scale database with 1000 molecules, and an FDA-approved drug.

To develop this chemoenzymatic synthesis planning tool, we used a deep learning model to design a score function named RTscore that can distinguish synthesis reactions from decomposition reactions and evaluate reaction effectiveness. In principle, RTscore could also be used separately to rank predicted synthesis routes, and RTscore_(ECREACT)_ was a specifically trained score for enzymatic reactions performing better than SCScore on an external enzymatic database. While our method focuses on selecting efficient reactions based on molecular transformations, the lack of reaction condition data in the USPTO and ECREACT databases may limit its practical applicability in some cases. Additional metrics, such as price, reaction conditions, and toxicity of chemicals, could be incorporated in the future for more comprehensive evaluations. In our hybrid synthesis search algorithm, RTscore was used to guide transitions between chemical and enzymatic reactions. This search method enabled the discovery of new enzymatic reactions while deepening the search tree to reach stock molecules efficiently. Since only the selected intermediate was searched for each route, RTscore also helped to save searching time.

The versatility of RTS-CESP was demonstrated in the combination with both ML-based and rule-based synthesis planning tools. In this work, five different tools were used to generate synthesis routes. The RXN4Chemistry (chemical) and RXN4Chemistry (enzymatic) employed a Molecular Transformer architecture and were continuously updated by the IBM researchers. The Aizynthfinder was trained with rules extracted from the Reaxys database and used MCTS as a multi-step search algorithm, making it a robust retrosynthesis tool. RetroBioCat used curated enzymatic reaction rules and has been verified by enzymatic cascades reported in literature, while BKMS extracted abundant enzymatic templates from four biological databases.

The robustness and reliability of RTS-CESP were validated by different tasks. For robustness, predicting synthesis routes for molecules in a large dataset could examine the competence of a synthesis planning tool in generating abundant transformations and reaching the commercially available molecules efficiently. Guided by RTscore, RTS-CESP predicted chemoenzymatic synthesis routes for more target molecules and suggested more shorter synthetic routes than ASKCOS. For reliability, it could be evaluated either using literature data or experimental validation. Other than reproducing and prioritizing the hybrid synthesis routes in our dataset with literature support, we carried out experiments to validate the synthesis route for the FDA-approved drug Alclofenac. The validated route is shorter than the literature-reported route, providing a promising synthesis option.

## ASSOCIATED CONTENT

## Data Sharing Agreement

The source code is available at https://github.com/Zhao-Group/RTS-CESP.

## Supporting Information

The following files are available free of charge.

Materials and methods, data cleaning, model development and evaluation, calculation parameters, experimental procedures (PDF)

## AUTHOR INFORMATION

## Corresponding Author

Huimin Zhao − NSF Molecular Maker Lab Institute, Department of Chemical and Biomolecular Engineering, Carl Woese Institute for Genomic Biology, Department of Chemistry, DOE Center for Advanced Bioenergy and Bioproducts Innovation, University of Illinois at Urbana-Champaign, Urbana, IL 61801, USA. orcid.org/0000-0002-9069-6739; Phone: (217) 333-2631; Fax: (217) 333-5052

## Authors

Hongxiang Li − NSF Molecular Maker Lab Institute, Department of Chemical and Biomolecular Engineering, Carl Woese Institute for Genomic Biology, Department of Chemistry, University of Illinois at Urbana-Champaign, Urbana, IL 61801, USA.

Xuan Liu − NSF Molecular Maker Lab Institute, Department of Chemical and Biomolecular Engineering, Carl Woese Institute for Genomic Biology, University of Illinois at Urbana-Champaign, Urbana, IL 61801, USA.

Guangde Jiang − Department of Chemical and Biomolecular Engineering, Carl Woese Institute for Genomic Biology, DOE Center for Advanced Bioenergy and Bioproducts Innovation, University of Illinois at Urbana-Champaign, Urbana, IL 61801, USA.

## Author Contributions

H.Z. coordinated the project. H.L. and H.Z. conceived the presented idea. H.L. conducted the computational and experimental studies. X.L. contributed to the benchmark study on the boutique database, while G.J. assisted in analyzing products in the experiment. H.L. and H.Z. wrote the manuscript with input from all authors.

## Notes

The authors declare no competing interests.

## ACKNOWLEDGMENT

We thank Dr. Haiyang Cui, Dr. Zhengyi Zhang, Dr. Maolin. Li and Zhenxiang Zhao, for useful suggestions in experiment.

